# Spatial variation in relative abundances of two butterfly species sharing both host plant and natural enemies

**DOI:** 10.1101/2022.11.23.517710

**Authors:** P. Colom, A. Traveset, M. R. Shaw, C. Stefanescu

## Abstract

The decline of insect populations is of great concern because they play an essential part in several services that are key for ecosystem functioning and human well-being. Therefore, full understanding of the processes and factors shaping spatial variation in insects is required for their effective conservation. Here, we study a system comprising two congeneric butterfly species (Brimstone *Gonepteryx rhamni* and Cleopatra *G. cleopatra*) that share both host plants and natural enemies and analyse whether biotic and/or abiotic factors explain their relative abundances. The two species coexist in continental Spain but not on a nearby archipelago, where only the Cleopatra occurs. The hypotheses tested were based on (H1) dispersal behaviour; (H2) apparent competition mediated via shared parasitoids; and (H3) environmental conditions (overwintering habitat availability, abundance of host plants and temperature). H1 explained differences in Brimstone abundance between climate regions on the mainland since in warmer summers populations increased in cooler areas but decreased in warmer areas. Cleopatra did not show the same pattern but was found to have twice the number of summer adults on one island than on the mainland. It is unlikely that H2 can explain this result because, although richer parasitoid communities were found on the mainland, larval mortality rates were similar. H3 was important in explaining variation in abundances between sites within each climate region even though similar environmental conditions were found on the mainland and on the islands. Our study demonstrates the complexity of any attempt to understand insect population dynamics in space due to the number of factors that are potentially involved. We argue thus that a more comprehensive approach taking into account landscape topography and resource connectivity on a broader scale is required to unravel the factors shaping the relative abundance of insects in island systems.

## INTRODUCTION

Insects represent more than half of the biodiversity on Earth and play a central role in trophic networks through many different types of biotic interactions and involvement in several processes as herbivory, detrivory, nutrient recycling or pollination (Chapman 1998). Consequently, the diversity of insects over space and time shapes ecological networks and ecosystem functioning. However, species occurrence and the population densities that determine the strength of biotic interactions are both relevant.

Insect population density, as in many other taxa, is regulated by biotic and abiotic factors: species experience different climatic conditions and occur in a variety of assemblages where the presence or absence of predators and competitors throughout their distributions drive abundances over space and time. However, when populations of the same species live under similar climatic conditions, differences in densities should be explained above all by differences in biotic interactions. In this regard, island biogeography theory states that species densities on islands may be higher than on the mainland due to competitive release resulting from the species impoverishment of island assemblages (Whittaker and Fernández-Palacios 2007). Nevertheless, competitive release in insect communities seems unlikely to occur because interspecific competition in phytophagous insects has been suggested (Lawton and Strong 1981, Shorrocks et al. 1984) and empirically shown to be rare (e.g. Kaplan and Denno 2007). Indirect interactions may play a more important role in structuring insect communities (Holt 1977). The sharing of natural enemies (i.e. predators or parasitoids) between species may account for the decrease in the abundance of some of these species when they coexist, the so-called ‘apparent competition’ (see a review of this concept in Holt and Bonsall 2017). Apparent competition in insect communities has been demonstrated to exist under controlled conditions in experimental settings (Bonsall and Hassell 1997), even though conclusive evidence is rare under natural conditions (but see e.g. Van Nouhuys and Hanski 2000, Frost et al. 2016, Audusseau et al. 2021).

Other factors such as the availability of suitable habitats and food resources should also be considered when attempting to unravel intraspecific density differences. These factors are especially relevant for species with strict ecological requirements such as monophagous insects or insects that only overwinter in particular habitats (Dennis et al. 2003, Dennis 2010). Moreover, a number of factors may interact and influence how the abundance of a species is distributed over space. For instance, the population dynamics of the small tortoiseshell *Aglais urticae* in the southern part of its range are driven by the interaction between aridity, host plant quality and larval parasitism (Stefanescu et al. 2022).

Here, we investigate two closely related butterfly species, *Gonepteryx rhamni* (Brimstone) and *G. cleopatra* (Cleopatra), that coexist on a mainland (Iberian Peninsula), of which one (Cleopatra) is also present on nearby continental islands (Balearic Islands). On the mainland, the two *Gonepteryx* species occupy different climatic niches, although their distribution broadly overlaps in the warmer area of the Brimstone’s range, which corresponds to the colder area of the Cleopatra’s distribution. Both these butterfly species use the same plants of the genus *Rhamnus* for larval development and are known to be parasited by specialist wasps on the mainland (Shaw et al. 2009); by contrast, no information was available about the parasitoid complexes occurring on the Balearic Islands. The complexity of the population dynamics of these two species increases when their highly dispersive behaviour is considered, and pre-overwintering and post-overwintering adults occur in different areas (Pollard and Hall 1980, Gutiérrez and Wilson 2014).

This work aims to identify the drivers shaping spatial variation in the relative abundance of herbivorous insects using a study system with a marked climatic gradient and data on species phenology and behaviour. In this regard, we first calculate and compare population densities of the two species in different biogeographic and climate regions, including two islands in the Balearic archipelago and several sites on the mainland, and consider separately post-overwintering and pre-overwintering adults. The predictive ability of abiotic and biotic factors to explain population densities in each flight period (pre- and post-overwintering) and climate region is tested. Then, taking into account the effect of these factors, we explore whether they are similar in magnitude in the two biogeographic regions. We also attempt to understand how important natural enemies (i.e. parasitoids) are to the variation in relative abundance through indirect interactions. Testing apparent competition in natural systems requires long-term studies tracking both hosts and parasitoid species over time and space, which is an inherently difficult challenge (Holt and Bonsall 2017). However, we were able to collect data for four years on parasitoid complexes and on their importance as a cause of mortality in *Gonepteryx* species on both the mainland and on the two nearby Balearic islands to test whether or not apparent competition between these two species could explain the high population density of Cleopatra on the islands. Long-term butterfly count data were used to test whether or not density dependence is higher when these two species coexist and whether or not it depends not only on the abundance of the same species but also on the abundance of the other co-occurring species as a possible sign of apparent competition. Finally, we discuss which explanations account best for the variation in relative abundance across the biogeographic and climate regions present in our study system.

## METHODS

### Study system

We analysed the coexistence of Brimstone and Cleopatra in a Mediterranean area of ca 36,900 km^2^ in which butterflies are systematically monitored within the framework of a citizen science project (www.catalanbms.org). The study area encompasses two biogeographic regions (islands: Mallorca and Menorca – Balearic Islands; mainland: Catalonia and Andorra – NE Iberian Peninsula) and three distinct climate regions: Mediterranean xeric, Mediterranean mesic and alpine-subalpine (Fig. S1). These climate regions were established using a threshold of 21GDD (number of hours per year in which a temperature of 21°C is exceeded) and broadly correspond to those classified by Metzger et al. (2013) as (1) alpine-subalpine (0–150 GDD); (2) mesic (between 150–400 GDD); and (3) xeric (≥400 GDD).

The Brimstone is widespread in Europe but at its southerly limit it is mainly restricted to upland and/or more humid areas. In our study area, it is found only sporadically in the arid region and is completely absent from the Balearic Islands. By contrast, the Cleopatra is mainly found in the Mediterranean Basin, where it is common in mesic and arid regions including the Balearic Islands and, as a previous study has shown, flies in very high population densities on the island of Menorca (Colom et al. 2019). Although the two species have different climatic niches (Settele et al. 2008), in our study area they often co-occur in the Mediterranean mesic zone, where they are amongst the commonest species. Both species occur in a wide range of habitats (Fig. S2) but depend on woodland for overwintering.

The Brimstone and Cleopatra are very closely related phylogenetically (Dapporto et al. 2022) and have similar phenologies and hostplant use (Vila et al. 2018). Although they overwinter in the adult stage, the Brimstone is strictly univoltine. The Cleopatra, on the other hand, has a more plastic phenology with a partially bivoltine life cycle in many years (albeit with a very low relative abundance in the second generation: see Fig. 1). In both species, mating occurs soon after the adult butterflies come out of hibernation in early spring (henceforth, overwintered adults). Oviposition is concentrated in spring and eggs are laid individually on the underside of the most tender leaves of the host plants. In the NE Iberian Peninsula, they both use *Rhamnus alaternus* as their main larval resource, which does not seem to be a limiting factor for their populations as it is frequent and abundant in Mediterranean habitats. In the xeric region, oviposition has also been recorded on *R. lycioides* an in Mallorca on the Balearic endemic *R. ludovici-salvatoris* (PC pers. obs.). In the alpine region, where *R. alaternus* does not occur, the main host plants of *G. rhamni* are *R. frangula*, which is also a frequent resource in northern Europe (Gutiérrez and Thomas 2000), and *R. alpina*. In general, the adults of the annual generation of both species emerge in June–July (henceforth, summer adults), although their flight periods vary depending on the climate region (Fig. 1).

**Figure 1.**
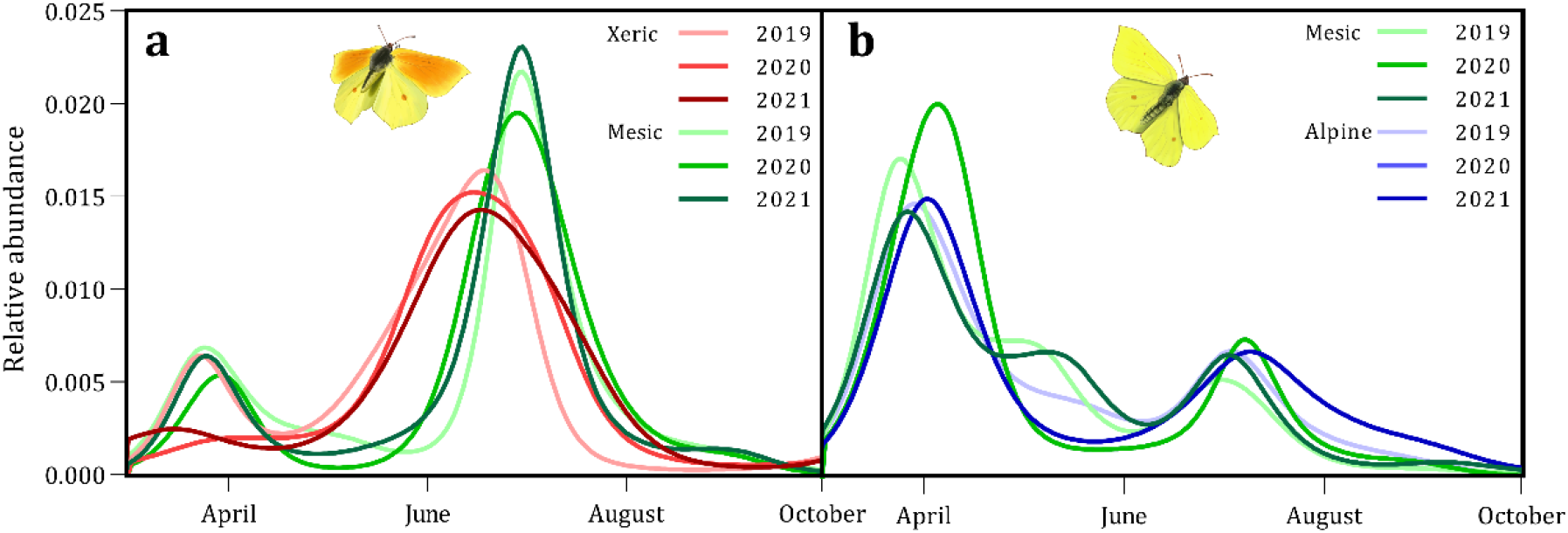
Flight curves estimated by GAM models using count data from 2019, 2020 and 2021 to illustrate the adult phenology of (a) Cleopatra and (b) Brimstone in each climate region. The first peak represents the overwintered adults and the second the summer adults. The relative abundances of each flight curve are standardized to add up to 1.

The Cleopatra and Brimstone potentially share solitary hymenopteran larval parasitoids specific to their genus: *Hyposoter rhodocerae* (Ichneumonidae: Campopleginae), *Cotesia gonopterygis* and *C. risilis* (Braconidae: Microgastrinae) (Shaw et al. 2009; Jubany and Stefanescu 2009). They are also attacked by other more generalist hymenopteran parasitoids, including egg *(Trichogramma cordubensis*: Trichogrammanidae; CS unpublished data) and pupal (*Pteromalus apum*: Pteromalidae; Shaw et al. 2009) parasitoids.

Both butterfly species are highly dispersive and several studies have provided evidence of seasonal migrations between habitats (Pollard and Hall 1980), including altitudinal migrations (Jubany and Stefanescu 2009, Gutiérrez and Wilson 2014). Gutiérrez and Wilson (2014) found that Brimstone adults in summer flew at sites that were on average 3°C cooler than their breeding sites. Their great dispersive behaviour explains their occurrence at sites where neither breed (Gutiérrez and Thomas 2000).

### Abundance data

Butterfly populations have been monitored at 205 sites in the whole study area since 1994, of which 139 were active in 2019–2021. At each site, adult butterflies were counted along fixed transects in a space 2.5 m on each side and 5 m in front of the observer (Pollard and Yates 1994). Abundance data were recorded weekly from March to September (i.e. 30 recording events per year, weather permitting). To estimate annual abundances, only male counts were used as they are much more easily identified than females to species level in the field. For each year we fitted flight curves for each climate region using generalized additive models (GAMs), following the approach of Schmucki et al. (2016). The GAMs fit bimodal curves that represent the post-overwintering and pre-overwintering flight periods of each species in each year and climate region (Fig. 1). This allowed us to estimate separately the total abundance of overwintering and summer adults in each population for each year.

### Larval sampling

We studied larval parasitism at several sites on Mallorca and Menorca and in Catalonia (including the xeric and mesic climate regions) to test the possibility of apparent competition between the two *Gonepteryx* species. From late March to early June over a period of four years (2019–2022), we collected 949 larvae from 14 sites (Mallorca: 6 sites; Menorca: 4 sites; Catalonia: 4 sites). Larvae were reared indoors using transparent plastic containers (155 × 105 × 45 mm) in groups of up to five individuals from the same sample. We recorded the larval instar for each collected individual. When a caterpillar was killed by a *Cotesia* parasitoid, we waited for the cocoon of the parasitoid to be fully formed and hardened before isolating it in a corked glass tube (80ml). When *Hyposoter rhodocerae* kills the host, it spins its cocoon within the host’s larval skin, after which we also isolated its structure in a corked glass tube. Once emerged, the adult parasitoids were preserved in pure ethanol. All parasitoids were identified by MRS.

The DNA sequencing of the butterfly larvae killed by *Cotesia* parasitoids on the mainland was carried out following standard protocols (Ivanova et al. 2006, deWaard et al. 2008, Hebert et al. 2013) at the Centre for Biodiversity Genomics, University of Guelph, Canada. All new sequences are deposited in BOLD, which are publicly available at dx.doi.org/10.5883/DS-GONCOLOM.

Parasitism rates were compared between regions and years with GLMs assuming a binomial distribution and logit link function.

### Environmental variables

To analyse the biotic and abiotic determinants of species abundance, we collected field data on habitat structure and host plant availability, and extracted climate data from an online source (Cornes et al. 2018). Between 2019 and 2021, plant communities (according to the CORINE land cover classification: Vigo et al. 2005) and cover of *Rhamnus* species were sampled at 84 random sites in the butterfly monitoring network. At each of these sites we then calculated the percentage cover of all types of forest communities as an estimate of habitat availability for overwintering adults. Coverage of host plants (i.e. m^2^ of *Rhamnus* sp.) along the 5-m wide butterfly transect was also calculated for the whole transect and standardized to a common area of 1000 m^2^ to allow transects to be compared.

We extracted daily maximum temperature data for the same subset of 84 sites at 0.1-degree resolution (ca 11 km in latitude) from the Copernicus Climate Change Service (Cornes et al. 2018). For each site and year, the maximum average temperatures in spring (March–May) and summer (June–August) were calculated given that – according to Gutiérrez and Wilson (2014) – these variables explain larval development and adult thermoregulation, and thus determine the distribution of the abundance of butterflies over space.

### Statistical analyses

#### Population density across biogeographic and climate regions

To test for differences in the density of populations between different climate and biogeographic regions we used data from 139 sites for which data were available for 2021. We used generalized linear models (GLMs) with a Gaussian distribution and a zero-inflation parameter (Brooks et al. 2017a). Abundance estimates were standardized by transect length and log-transformed to reach normality. We performed four models in which the response variable was the abundance of each flight period (overwintering and summer adults) of each species (Brimstone and Cleopatra). The independent variable was a categorical factor of four levels (Mallorca, Menorca, Cat. xeric and Cat. mesic) for the models of Cleopatra, and two levels (Cat. mesic and Cat. alpine-subalpine) for the models of Brimstone. We repeated the analyses for both species using generalized linear mixed models (GLMMs) for a data set of 205 sites and 28 years (1994–2021), with year as a random effect.

#### Environmental determinants of population density

The predictive power of the three selected environmental variables (forest cover, host plant cover and spring/summer maximum average temperatures) on the abundance of each generation of each species in each climate region was tested (eight models in total). Predictor variables were rescaled to values ranging from 0 to 1. We used data from the 84 sites for which data on both habitat and butterfly abundance were available.

The response variable was the mean standardized abundance for 2019–2021. Because the data were extremely unbalanced to 0, GLMs were performed with a Tweedie distribution that best fitted the models (Brooks et al. 2017a).

The environmental variables were compared between biogeographic and climate regions using the same structure as the analyses of population densities (see above).

#### Density dependence models

A recent study has shown that density dependence in our study region plays an important role in the population dynamics of most butterfly species including the Brimstone and Cleopatra (Ubach et al. 2022). Density dependence is probably related to the impact of natural enemies, particularly parasitoids, which have been shown by many butterfly studies carried out in the region to represent a major cause of mortality (e.g. Shaw et al. 2009, Stefanescu et al. 2022). Because Cleopatra and Brimstone potentially share the same specialist parasitoids in our region (see Results), we investigated the possibility that apparent competition was mediated by shared parasitoids on the mainland (Bonsall and Hassell 1997) where the two butterfly species co-occur (Fig. S1). Under apparent competition, we expected that the inter-annual population growth of one species would depend not only on its own density in the previous year but also on the density of the other species (henceforth, inter-specific density dependence). To test this hypothesis, we used sites with data from at least 10 consecutive years in which both species coexist (i.e. the Mediterranean mesic region; Fig. S1; n = 41). For each species *i*, the response variable of the models was the inter-annual population growth calculated as the difference between the log-transformed abundance in the current year (*t*) and in the previous year (*t-1*). The predictors of the models were (a) the abundance of the species *i* in *t-1*; (b) the abundance of the species *j* in *t-1*; and (c) the sum of the abundance of the species *i* and *j* in *t-1*. Because predictor *c* was highly correlated to predictors *a* and *b*, it was included in a model as the only predictor. For each species, GLMMs were carried out with all the combinations of the predictors and we then selected the best model based on AIC. For all the models, a Gaussian distribution including a zero-inflation parameter was used, with site controlled as a random effect. Finally, we compared the magnitude of the effect of the density dependence of species *i* between the region in which both species coexist and the regions in which they do not.

All analyses were carried out using R v. 4.2.1 (R Core Team 2020) and the following packages: *climateExtract* to extract and manipulate ECAD climate data (Schmucki 2022), *rbms* to fit the flight curves (Schmucki et al. 2022), *glmmTMB* to conduct GLMs and GLMMs (Brooks et al. 2017b), and *MuMIn* (Barton 2020) for model comparison.

## RESULTS

### Predictors of population density

The importance of each of the different environmental factors in predicting butterfly abundance depended on the climate region (Table 1). The availability of host plant resources and forest habitat explained the abundance of Cleopatra in the cooler region (mesic), while the maximum temperature constrained its abundance in the warmer region (xeric); nevertheless, this was only significant for the overwintered adults. Similarly, the abundance of Brimstone summer adults was constrained by the maximum temperature in the warmer region of its distribution (mesic), while forest cover was also found to be important for the overwintered population. In both species, a higher abundance of overwintered adults occurred at sites with greater forest cover. In the alpine-subalpine and cooler climate region, forest cover predicted the abundance of both the overwintered and summer adult Brimstones, while temperature had a significant positive effect on summer adults. Unlike the Cleopatra, the availability of host plant resources did not predict the abundance of the Brimstone.

**Table 1.**
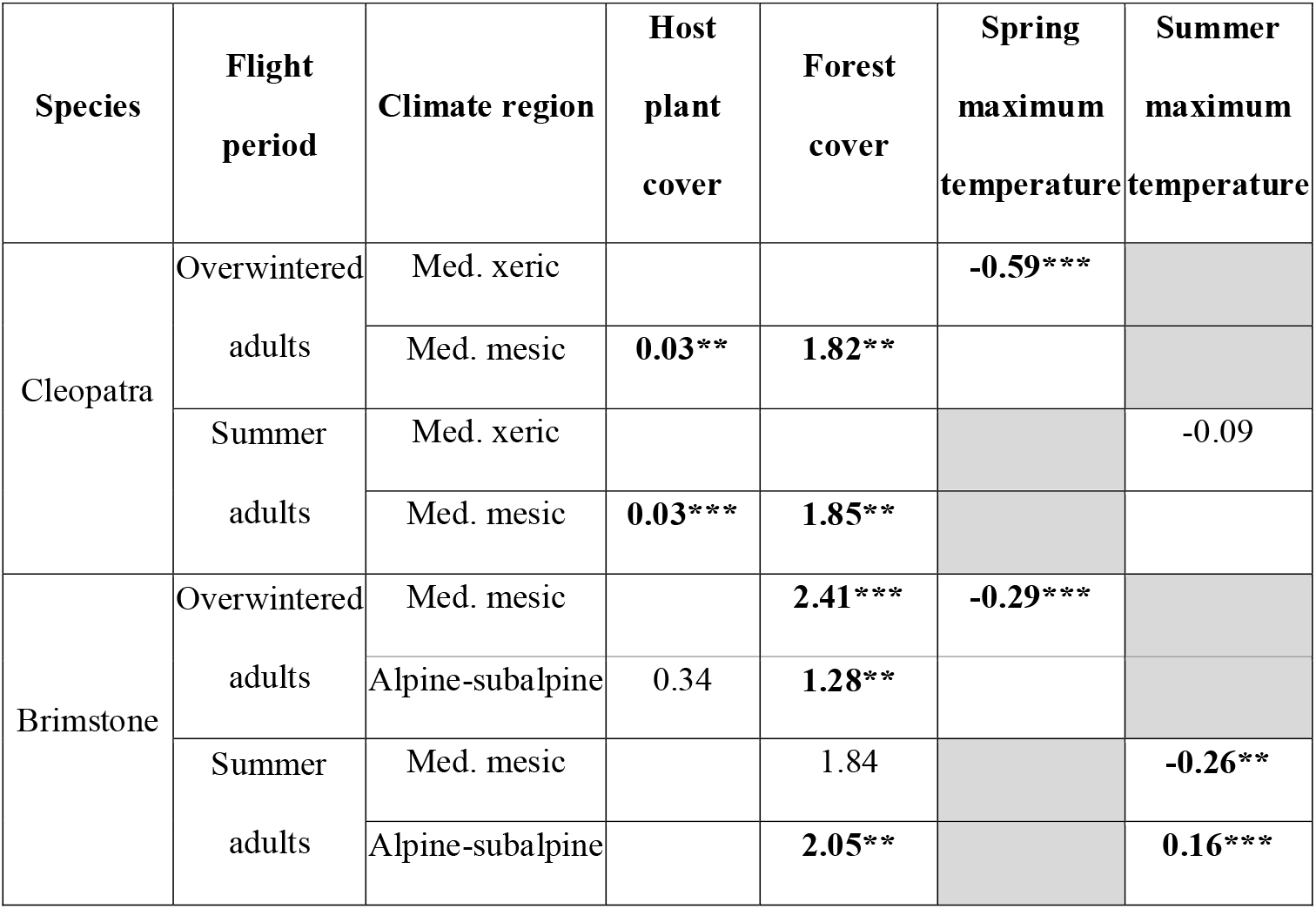
Results of the GLMs testing the effect of the environmental predictors on butterfly abundance. Values of the coefficients for the variables included in the best-fit model based on the AIC are shown. Values with a significant effect are shown in bold with asterisks (* *P*< 0.1; ** *P* < 0.05; *** *P* < 0.01).

Two biotic factors (forest and host plant cover) were well represented throughout all the biogeographic and climate regions (Fig. 2a,b). However, no host plants were found in the alpine-subalpine sites, while the mainland sites in the xeric regions had significantly higher levels of host plant cover in comparison with selected sites on Mallorca. Spring and summer average temperatures were significantly lower in the alpine-subalpine region (Fig. 2c,d).

**Figure 2.**
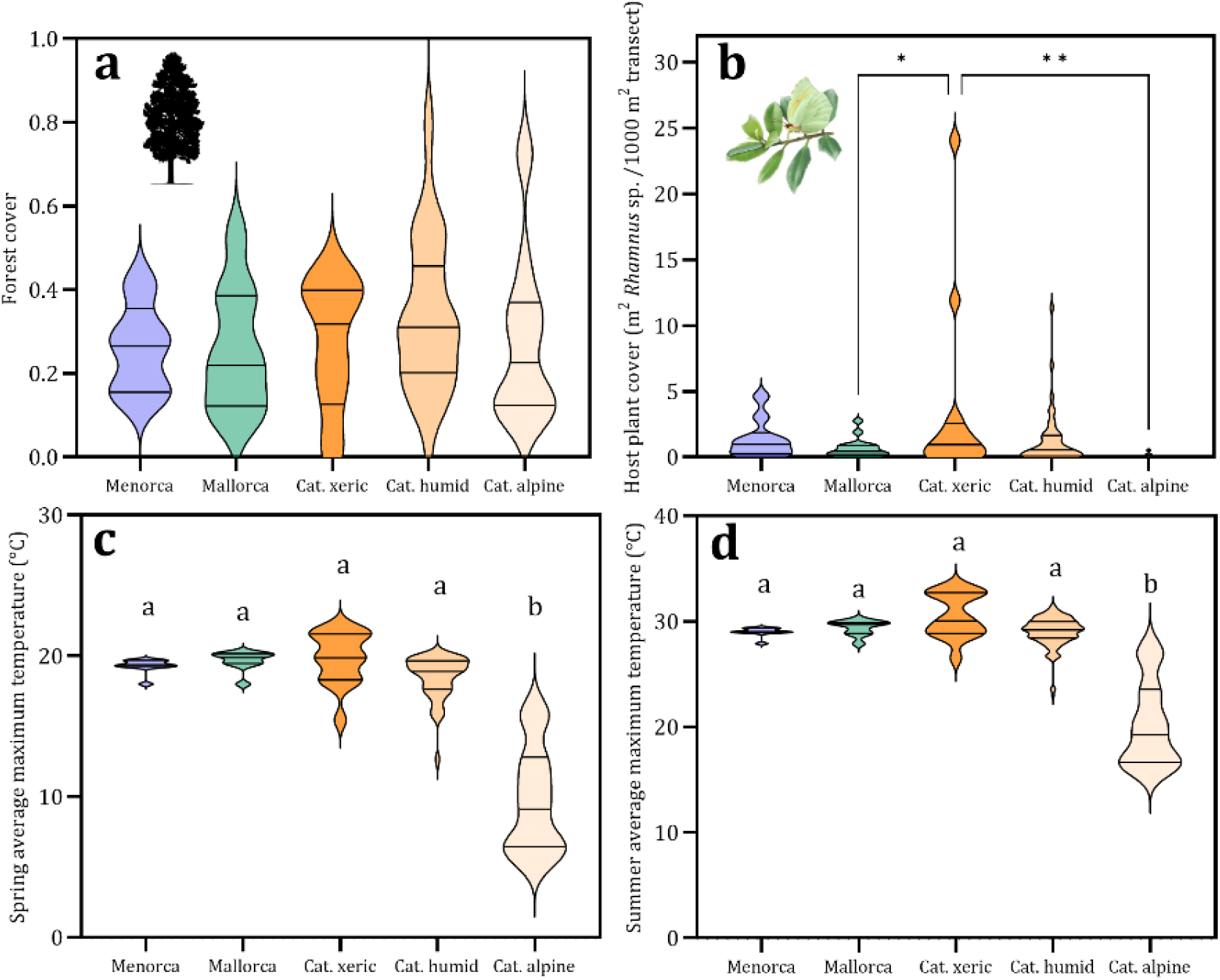
Comparison of the environmental variables between biogeographic and climate regions. (a) Total percentage (0–1) of cover of woodland habitats along the butterfly transects. (b) Host plant cover (*Rhamnus* sp.) standardized per 1000 m of transect length. (c) Spring (March–May) average maximum temperature. (d) Summer (June–August) average maximum temperature. Significant differences between the groups are shown with different letters or with asterisks (* *P* < 0.05; ** *P* < 0.01).

### Population density across biogeographic and climate regions

Overwintered individuals of both species had similar population densities throughout the study area regardless of biogeography or climate. By contrast, we found significant differences between regions for summer adults of both species. Cleopatra had significantly higher population densities on Menorca (0.52 ± 0.36) than on both Mallorca (0.2 ± 0.18) and the two mainland climate regions (xeric: 0.15 ± 0.24; mesic: 0.28 ± 0.22). On the mainland, the Brimstone was more abundant in the cooler (alpine-subalpine: 0.18 ± 0.18) than in the warmer regions (mesic: 0.09 ± 0.1). Population densities for each combination of species-flight period-region are shown in Figure 3. The same qualitative results were obtained using the GLMMs (see Methods).

**Figure 3.**
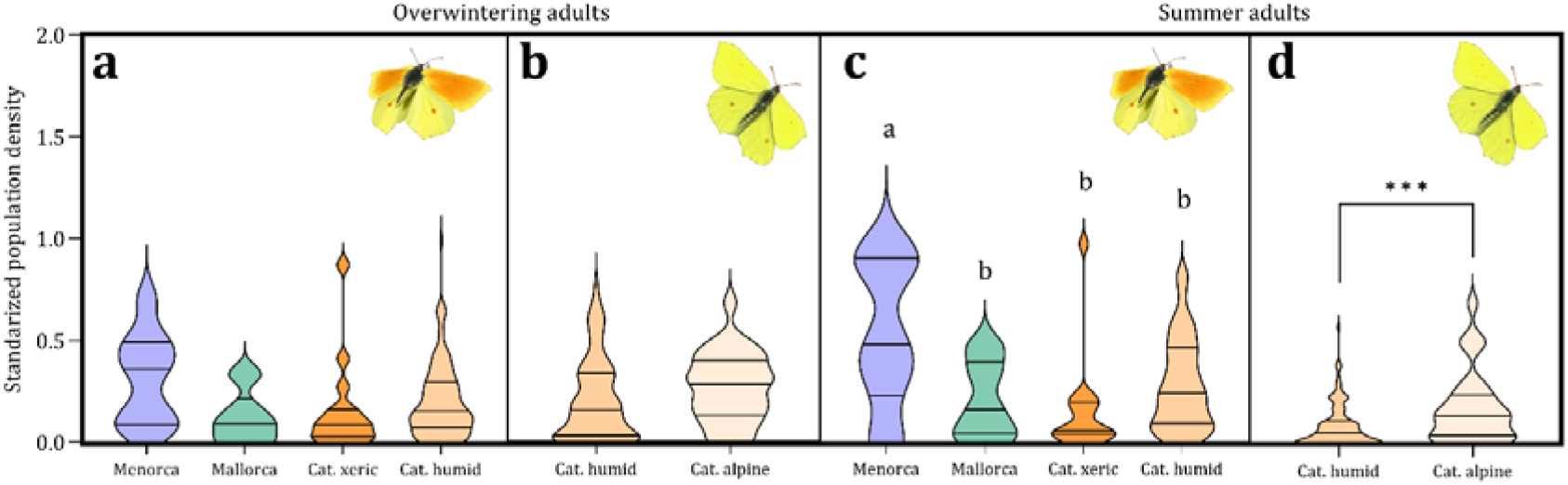
Comparison of populations densities between biogeographic and climate regions. Overwintered adults of (a) Cleopatra and (b) Brimstone; summer adults of (c) Cleopatra and (d) Brimstone. Letters and asterisks show significant differences between groups. Population density estimates were standardized by transect length and log-transformed to reach normality.

### Larval parasitism

During the study we collected 949 larvae from Menorca, Mallorca and Catalonia (including sites in the xeric and mesic climate regions): 103 larvae from Menorca in 2019, 412 from Mallorca (2019: 152; 2020: 122; 2021: 110; 2022: 28) and 434 from Catalonia (2021: 144; 2022: 290). However, 35% of the larvae died during rearing for unknown reasons and were excluded from the calculations of parasitism rates. Parasitism was an important source of mortality in *Gonepteryx* larvae (Fig. 4). On the islands, the percentage of larvae killed by parasitoids ranged from 15% to 45% on Mallorca, and was 28% in Menorca in 2019. In Catalonia, where total parasitism rates were lower than on the islands, a total of 9% of larvae died from parasitism in both 2021 and 2022.

**Figure 4.**
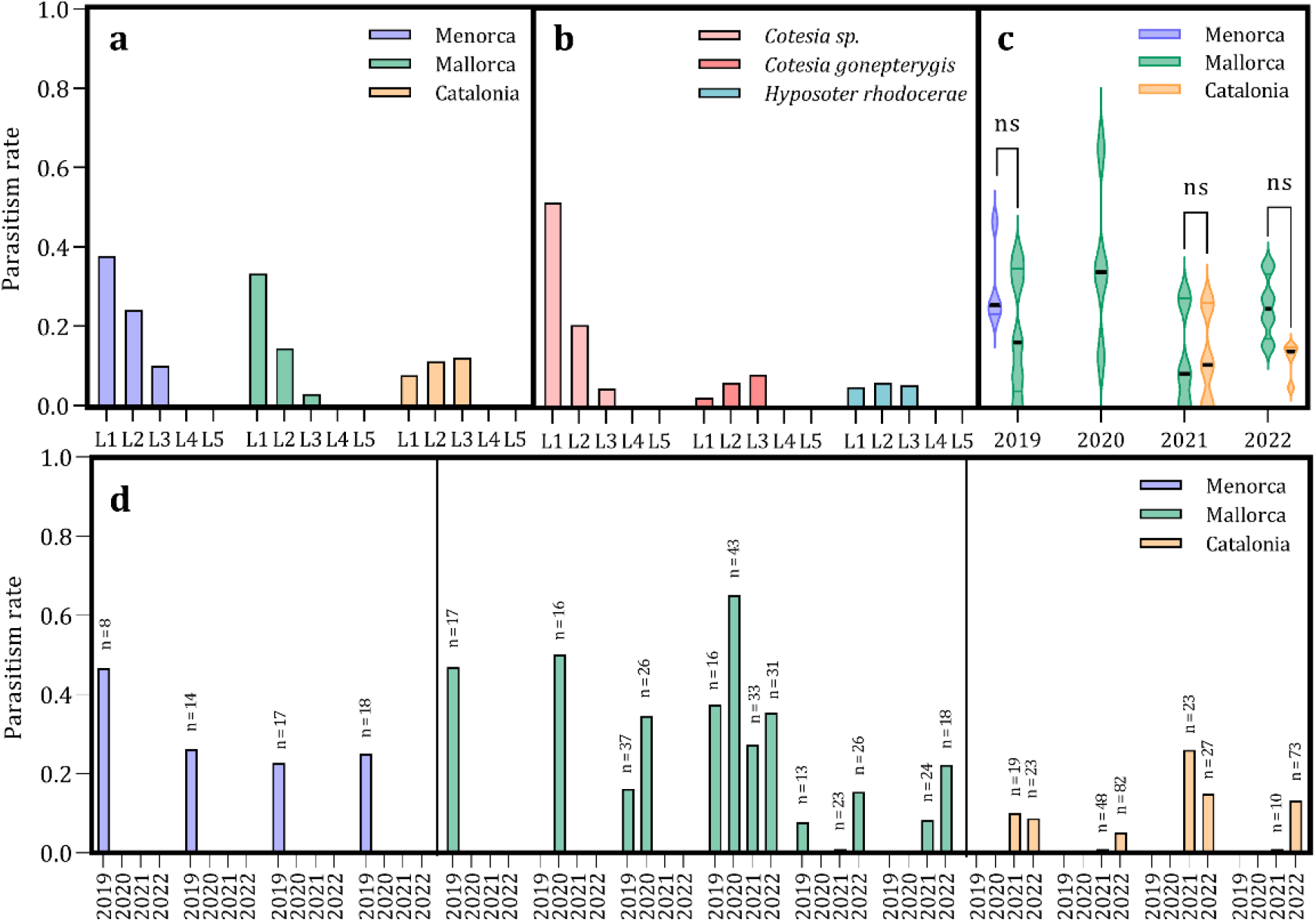
Parasitism rates in *Gonepteryx* larvae. (a) Total regional parasitism rates by larval instar (L1-L5) at the moment they were collected in the field. (b) Total parasitism rates for each parasitoid species by larval instar. (c) Comparison of local parasitism rates (site level) between regions in the same year. (d) Variability of parasitism rates between sites and years. Each bar represents the parasitism rate for a specific site and year.

Interestingly, different parasitoid species occurred on the mainland and the islands. At mainland sites, two species known to be specialist parasitoids of the genus *Gonepteryx* (Shaw et al. 2009), the braconid and the ichneumonid wasps *Cotesia gonopterygis* and *Hyposoter rhodocerae,* were recorded. All adults of *C. gonopterygis* emerged from *G. rhamni* larvae (n = 12), most of them from the sample taken from the coolest sampled site on the mainland (in El Montseny mountains), while only two individuals were recorded from the warmest site (Argentona). We could not identify the hosts of *H. rhodocerae* to species level but they were found at all except the coolest site, including a site where no larvae of *G. rhamni* was found (Sant Quintí, see Supplementary Figure S4). Surprisingly, *Cotesia risilis*, a fairly specialist parasitoid of *Gonepteryx* species in the study area (Jubany and Stefanescu, 2009), did not appear at any site. On the islands, the Cleopatra was only attacked by one as yet undescribed endemic parasitoid of the genus *Cotesia* (Shaw and Colom in prep.).

Parasitism was concentrated in the first three larval instars, although the distribution of parasitized larvae by instars showed differences between the mainland and the islands (Fig. 4a). On the islands, the endemic *Cotesia* seems mainly to parasite larvae in their first instar because parasitism rates decreased steadily from the first to the third instar at the time of collection. On the mainland, on the other hand, parasitism rates were more homogenously distributed among the larvae collected throughout the three first instars (Fig. 4b). The parasitoids generally killed the larvae in the third instar. No statistical differences in parasitism rates were detected between regions (Fig 4c). Fig. 4d illustrates the variability in parasitism rates between sites in the same region between years.

### Density dependence

Both the density of the same species and the sum of the densities of the two species had significantly negative effects on the inter-annual population growth of the Brimstone and Cleopatra (Supplementary Table S1). Inter-annual population growth was best predicted by the density of the same species in *t-1* than by the sum of the densities of the two species (Cleopatra: dAIC = 41; Brimstone: dAIC = 118.9). Models only including the inter-specific density effect also had a significant effect but of lower magnitude (Cleopatra: −0.5 vs −0.07; Brimstone: −0.62 vs −0.06) and had the highest AIC values (Cleopatra: dAIC =88.7; Brimstone: dAIC = 132.8).

The strength of the density dependence varied significantly between climate regions but not between biogeographic regions (Supplementary Table S2). Density dependence was greater in the mesic than in the xeric region for Cleopatra, while this same effect was greater in the alpine-subalpine than in the mesic region for Brimstone (Fig. 5).

**Figure 5.**
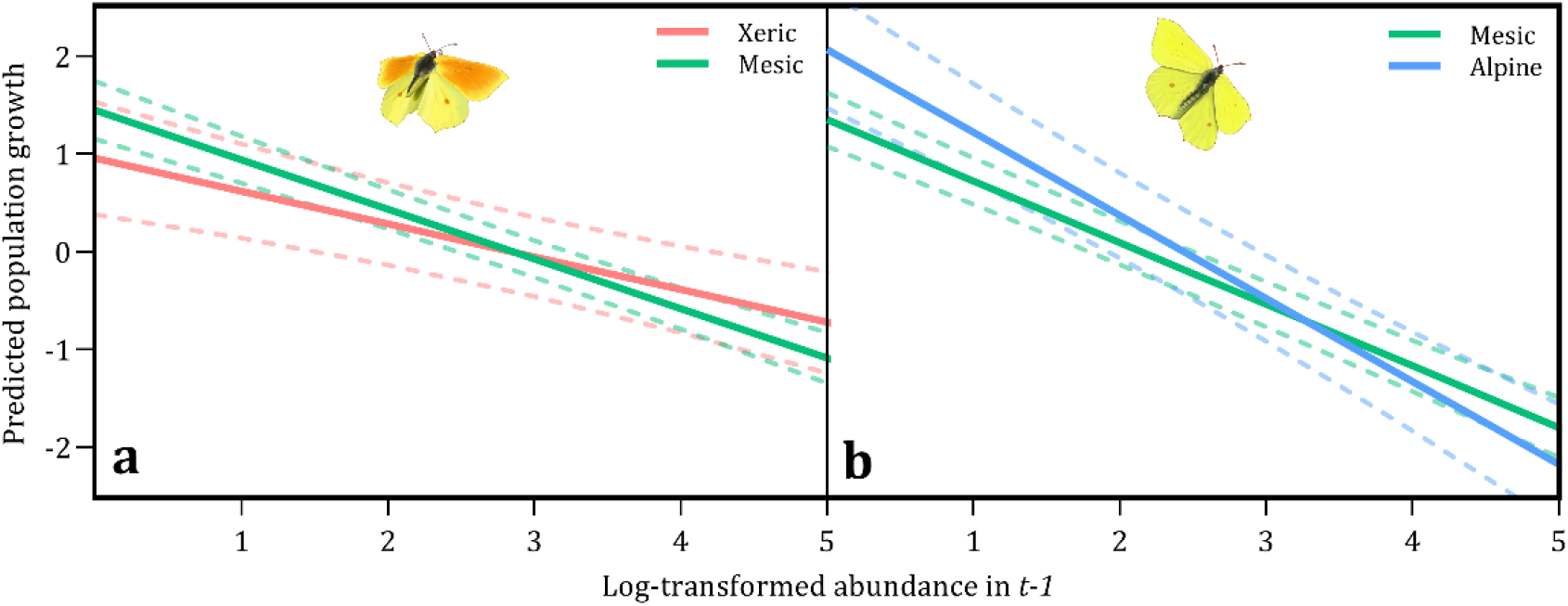
Predictor effect plots of density dependence differences between climate regions. Density dependence was measured as the relationship between population growth and abundance in *t-1*. (a) Significant differences between the density dependent effect on Cleopatra in the xeric and mesic Mediterranean climate regions. (b) Significant differences between the density dependent effect on Brimstone in the Mediterranean mesic and alpine-subalpine climate regions. Lines depict model-predicted relationships between population growth and abundance in *t-1*. Dashed lines represent the 95% confidence intervals.

## DISCUSSION

Observational and experimental studies have shown that insect populations fluctuate strongly over space and time according to differences in environmental conditions, resources and biotic interactions. All these factors that have been related potentially to intraspecific variability in insect population densities have been explored in this work.

In a previous study we found evidence for greater population densities in the Cleopatra in Menorca than in their mainland counterparts (Colom et al. 2019). In the current work, we improved the complexity and reliability of the measures of these relative abundances. First, the number of study sites was substantially greater and Mallorca was included in the study system as an island with habitats and species assemblages similar to those found on Menorca. Second, mainland sites were clustered by climate region and, finally, because of the dispersal behaviour of adult butterflies, pre-overwintering and post-overwintering abundances were differentiated.

This new approach confirms the exceptional population densities of Cleopatra in Menorca, where densities are higher than in all mainland climate regions. Although the highest densities of post-overwintered adults were recorded on Menorca, differences from the other regions become statistically significant in summer with the emergence of the annual generation. For this generation, the mean density on Menorca is twice the mean density in the other regions (Fig. 3c). However, the numbers of Cleopatra recorded on Mallorca were closer to those observed on the mainland than to those on Menorca. We further detected important differences in the abundance of Brimstone summer adults between the two climate regions in which this species occurs on the mainland.

Below we discuss the role of the different factors explaining variation in the relative abundance of the two study species.

### Climate-driven dispersal behaviour

The greater abundances in the alpine-subalpine than in the Mediterranean mesic region of Brimstone summer adults (Fig. 3d) can be easily explained by the dispersal of adults to cooler sites in summer (Gutiérrez and Wilson 2014). The fact that host plants were not recorded at the alpine-subalpine study sites supports the idea that butterflies present at these sites in summer came from other areas. Indeed, our results suggest that higher temperatures increase the rates of adult dispersal from warmer to cooler sites, which means that the differences between these sites are likely to increase with future global warming (Table 1). On the other hand, no significant differences were found between Cleopatra population densities in xeric and mesic regions. This does not imply that summer adults of Cleopatra do not perform similar movements to those of the Brimstone; nevertheless, we may failed to find significant differences between regions simply because the differences in maximum temperatures between the xeric and mesic regions were not as strong as between the alpine and mesic regions (Fig. 2d). Moreover, the significant negative effect of maximum temperature on overwintered adult Cleopatras in the xeric region suggests that a major dispersal of individuals to cooler sites does occur in warmer years.

The movements of summer adults to cooler sites could be influenced by the availability of nectar resources. However, Gutiérrez and Wilson (2014) found no support for the resource availability hypothesis (based on flower abundance) as an explanation for the summer altitudinal migration of *G. rhamni* in central Spain. Migration in the Painted Lady *Vanessa cardui* has been discussed partly as an evolutionary response to parasitism pressure (Stefanescu et al. 2012), although this hypothesis does not apply to the dispersal movements of *Gonepteryx* adults because the vast majority of individuals do not reproduce in summer.

### Variation in parasitism pressure and apparent competition

The data obtained from larval sampling do not support the hypothesis that indirect interactions between Brimstone and Cleopatra on the mainland lead to lower abundances than on the islands since, first, compared to the mainland regions Cleopatra populations were more abundant on Menorca but not on Mallorca, and, second, because larval parasitism rates were not higher on the mainland than at the island sites. Indeed, although only one parasitoid species was found on the islands, it caused similar mortality rates to those provoked by the two parasitoid species combined on the mainland (Fig. 4). The endemic *Cotesia* species was found at all island sites and was found to have a great capacity to parasitize Cleopatras in a wide range of habitats and environmental conditions. On the other hand, our data suggest that the two parasitoids found on the mainland have a low niche overlap as they only co-occurred in one of the four sampled sites (Fig. S3). Specialist parasitoids tend to vary their use of temporal and spatial resources to facilitate coexistence (Hood et al. 2021), so more natural enemies on the mainland do not necessary imply higher levels of butterfly larval mortality. The low niche overlap between the two parasitoids on the mainland can be explained by climate, habitat and/or host preferences, differences that may have originated as a result of a phenomenon of competitive exclusion. Our findings suggest that *Cotesia gonopterygis* occurs at colder sites and *Hyposoter rhodocerae* at warmer sites, although *H. rhodocerae* at least is known to be a common parasitoid of *G. rhamni* in colder regions such as the UK (MRS pers. obs.). The fact that all *C. gonopterygis* individuals emerged from Brimstone larvae suggests that it specializes on this species; yet, *H. rhodocerae* was found at a site where we found no Brimstone larvae, which suggests that it can use both *Gonepteryx* species as a host. Therefore, although apparent competition could still be mediated by *H. rhodocerae*, surprisingly we found no *H. rhodocerae* at the site with a more equal representation of the larvae of both species (El Montseny, see Fig. S4).

Apparent competition between *Gonepteryx* species seems even more unlikely if we take into account the fact that larval coexistence was unexpectedly low despite the high adult overlap (Fig. S4). While Brimstone adults were common at three of the four sampled mainland sites, only at the coolest sites were the larvae of this species common too. Therefore, our results indicate that, although adults of both *Gonepteryx* species coexist over a vast area, there is a great segregation in the breeding areas that limits the possibility of indirect effects. This suggests that these two closely related species have differentiated their climatic niches more than might be expected given the spatial distribution of adults alone, and that this could be the result of an evolutionary process promoting coexistence (Duyck et al. 2006).

An indirect approach using the density dependence models also failed to give support for the hypothesis of apparent competition. We expected that if significant indirect effects occurred, the sum of the densities of both species in the previous year (*t-1*) would govern the inter-annual population growth of their populations. However, the density of the same species in *t-1* was the variable that best explained the inter-annual population growth (Supplementary Table S1). The only result supporting the apparent competition hypothesis were the higher levels of density dependence in the Cleopatra populations in the region where this species co-exists with the Brimstone than in the warmer region (Fig. 5a). Nevertheless, the opposite was found in Brimstone populations because the density dependence effect was higher in the region where the Cleopatra is not present (Fig. 5b). However, factors other than parasitism may account for density dependence processes in butterfly populations dynamics (Stiling 1988, Dooley et al. 2013).

### Overwintering habitat and abundance of larval resources

The availability of host plants and overwintering habitat play an important role in explaining the spatial heterogeneity of butterfly abundances (Yamamoto et al. 2007, Curtis et al. 2015, Flockhart et al. 2015). Host plant and forest cover significantly influenced the abundance of overwintering and summer Cleopatra adults suggesting that larval trophic resources and overwintering habitat are both important. By contrast, Brimstones at cooler sites were more limited by overwintering habitat than by host plant availability.

Differences in forest and host plant cover are unlikely to be large between the mainland and the islands and between the two islands, at least at the study sites (Fig. 2a,b). Therefore, the high densities of Cleopatra in Menorca do not appear to be due to increased resource abundance or habitat cover on this island. However, Cleopatra adults can perform long movements and our data set is restricted to the transect level. Thus, data on host plant and habitat cover over wider areas are needed to establish whether or not differences in summer populations between Menorca and the other regions are attributable to differences in environmental variables at larger scales.

### Alternative factors shaping spatial variation in butterfly relative abundance

Taking into account all these results, it is still difficult to find a clear explanation for the much higher population densities of Cleopatra on Menorca. Differences between islands could be attributable to island geography (size and distance from the mainland), regional conditions or island heterogeneity (e.g. topographic heterogeneity) (Dennis and Hardy 2018). Menorca is a small island (area: 695.7 km^2^ vs Mallorca: 3640.1 km^2^) with a very flat landscape (highest elevation: 385 m vs Mallorca: 1436 m), which means that there are fewer opportunities for summer adults to disperse to cooler sites than on Mallorca or the mainland (but see Colom et al. 2021). If dispersion is limited, summer individuals may be forced to spend more time flying at the same sites, thereby increasing their detectability relative to the sites in the other regions. Hence, the higher numbers recorded in Menorcan transects may in fact result from behavioural differences rather than real differences in population density.

Another hypothesis is that Menorca may have been less affected by land-use changes, urbanization and the loss of habitat heterogeneity over the past two decades than either Mallorca or the mainland given that it has been a Biosphere reserve since 1993. In spite of the high mobility of Cleopatra adults, capable of reaching suitable distant habitats and food resources, landscape structure may still play a role and enhance on Menorca the connectivity between resources and their exploitation and, consequently, maintain larger butterfly populations.

Overall, our work illustrates the complexity of attempting to disentangle the processes shaping the abundance of insects over space due to the many different factors that are potentially involved. It also highlights how important it is for biogeographic and macroecological studies to be able to count on good knowledge of the ecology of the studied species, as well as interspecific interactions of various kinds, for their analyses.

## ACKNOWLEDGEMENTS

We are very grateful to all the CBMS volunteers who gathered the butterfly data. We also wish to thank Andreu Ubach and Ferran Páramo for helping with data management, and Roger Vila for assisting us with the DNA analyses. The CBMS is funded by the Departament d’Acció Climàtica, Alimentació i Agenda Rural de la Generalitat de Catalunya, the Diputació de Barcelona and the Andorran Government (via BMSAnd project). P.C. is funded by a PhD fellowship financed by the Govern de les Illes Balears (grant no. FPI-CAIB-2018) within the DEPICT research project (grant no. PID2020-114324GB-C2) funded by the Spanish MCIU Ministry to A.T.

## DATA AVAILABILITY STATEMENT

We will archive all data with the Figshare repository.

## References

Audusseau, H., N. Ryrholm, C. Stefanescu, S. Tharel, C. Jansson, L. Champeaux, M. R. Shaw, C. Raper, O. T. Lewis, N. Janz, and R. Schmucki. 2021. Rewiring of interactions in a changing environment: nettle-feeding butterflies and their parasitoids. Oikos 130:624–636.

Barton, K. 2020. MuMIn: multi-model inference. R package version 1.43.17.

Bonsall, M. B., and M. P. Hassell. 1997. Apparent competition structures ecological assemblages. Nature 388:371–373.

Brooks, M. E., K. Kristensen, and K. J. Van Benthem. 2017a. Modeling zero-inflated count data with glmmTMB:1–14.

Brooks, M., K. Kristensen, K. J. van Bentham, A. Magnusson, C. W. Berg, A. Nielsen, H. J. Skaug, M. Maechler, and B. M. Bolker. 2017b. Package ‘ glmmTMB ’ R topics documented□: Page The R Journal.

Chapman, R. F. 1998. The insects: structure and function. Cambridge University Press.

Colom, P., D. Carreras, and C. Stefanescu. 2019. Long-term monitoring of Menorcan butterfly populations reveals widespread insular biogeographical patterns and negative trends. Biodiversity and Conservation 28:1837–1851.

Colom, P., A. Traveset, D. Carreras, and C. Stefanescu. 2021. Spatio-temporal responses of butterflies to global warming on a Mediterranean island over two decades. Ecological Entomology 46:262–272.

Cornes, R. C., G. van der Schrier, E. J. M. van den Besselaar, and P. D. Jones. 2018. An Ensemble Version of the E-OBS Temperature and Precipitation Data Sets. Journal of Geophysical Research: Atmospheres 123:9391–9409.

Curtis, R. J., T. M. Brereton, R. L. H. Dennis, C. Carbone, and N. J. B. Isaac. 2015. Butterfly abundance is determined by food availability and is mediated by species traits. Journal of Applied Ecology 52:1676–1684.

Dapporto, L., M. Menchetti, R. Vodă, C. Corbella, S. Cuvelier, I. Djemadi, M. Gascoigne□Pees, J. C. Hinojosa, N. T. Lam, M. Serracanta, G. Talavera, V. Dincă, and R. Vila. 2022. The atlas of mitochondrial genetic diversity for Western Palaearctic butterflies. Global Ecology and Biogeography: 1–7.

Dennis, R. L. H. 2010. A Resource-based Habitat View for Conservation. Butterflies in the British Landscape. Wiley-Blackwell, Oxford.

Dennis, R. L. H., and P. B. Hardy. 2018. British and Irish Butterflies: an Island perspective. CABI. Wallingford, UK.

Dennis, R. L. H., T. G. Shreeve, and H. Van Dyck. 2003. Towards a Functional Resource-Based Concept for Habitat□: A Butterfly Biology Viewpoint. Oikos 102:417–426.

deWaard, J. R., N. V. Ivanova, and P. D. N. Hebert. 2008. Assembling DNA barcodes: analytical protocols. Page Cristofre M, ed. Methods in molecular biology: environmental genetics. Totowa: Humana Press.

Dooley, C. A., M. B. Bonsall, T. Brereton, and T. Oliver. 2013. Spatial variation in the magnitude and functional form of density-dependent processes on the large skipper butterfly Ochlodes sylvanus. Ecological Entomology 38:608–616.

Duyck, P. F., P. David, and S. Quilici. 2006. Climatic niche partitioning following successive invasions by fruit flies in La Réunion. Journal of Animal Ecology 75:518–526.

Flockhart, D. T. T., J. B. Pichancourt, D. R. Norris, and T. G. Martin. 2015. Unravelling the annual cycle in a migratory animal: Breeding-season habitat loss drives population declines of monarch butterflies. Journal of Animal Ecology 84:155–165.

Frost, C. M., G. Peralta, T. A. Rand, R. K. Didham, A. Varsani, and J. M. Tylianakis. 2016. Apparent competition drives community-wide parasitism rates and changes in host abundance across ecosystem boundaries. Nature Communications 7:1–12.

Gutiérrez, D., and C. D. Thomas. 2000. Marginal range expansion in a host-limited butterfly species Gonepteryx rhamni. Ecological Entomology 25:165–170.

Gutiérrez, D., and R. J. Wilson. 2014. Climate conditions and resource availability drive return elevational migrations in a single-brooded insect. Oecologia 175:861–873.

Hebert, P. D. N., J. R. DeWaard, E. V. Zakharov, S. W. J. Prosser, J. E. Sones, J. T. A. McKeown, B. Mantle, and J. La Salle. 2013. A DNA “Barcode Blitz”: Rapid Digitization and Sequencing of a Natural History Collection. PLoS ONE 8:e68535.

Holt, R. D. 1977. Predation, apparent competition, and the structure of prey communities. Theoretical population biology 12:197–229.

Holt, R. D., and M. B. Bonsall. 2017. Apparent Competition. Annual Review of Ecology, Evolution, and Systematics 48:447–471.

Hood, G. R., D. Blankinship, M. M. Doellman, and J. L. Feder. 2021. Temporal resource partitioning mitigates interspecific competition and promotes coexistence among insect parasites. Biological Reviews 96:1969–1988.

Ivanova, N. V., J. R. Dewaard, and P. D. N. Hebert. 2006. An inexpensive, automation-friendly protocol for recovering high-quality DNA. Molecular Ecology Notes 6:998–1002.

Jubany, J., and C. Stefanescu. 2009. Gonepteryx rhamni i G.cleopatra, un toc de groc que marca el final de l’hivern. Cynthia, butlletí del Butterfly Monitoring Scheme a Catalunya 8:18–22.

Kaplan, I., and R. F. Denno. 2007. Interspecific interactions in phytophagous insects revisited□: a quantitative assessment of competition theory. Ecology Letters 10:977–994.

Lawton, J. H., and D. R. Strong. 1981. Community patterns and competition in folivorous insects. The American Naturalist 118:317–338.

Van Nouhuys, S., and I. Hanski. 2000. Apparent competition between parasitoids mediated by a shared hyperparasitoid. Ecology Letters 3:82–84.

Pollard, E., and M. L. Hall. 1980. Possible movement of Gonepteryx rhamni (L.)(Lepidoptera: Pieridae) between hibernating and breeding areas. Entomologist’s gazette 31:217–220.

Pollard, E., and T. J. Yates. 1994. Monitoring butterflies for ecology and conservation: the British butterfly monitoring scheme. Springer Science & Business Media.

R Core Team. 2020. R: A Language and Environment for Statistical Computing. R Foundation for Statistical Computing, Vienna, Austria.

Schmucki, R. 2022. climateExtract: Extract and manipulate daily gridded observational dataset of European climate (E-OBS) provided by ECA&D. R package version 1.25.

Schmucki, R., C. A. Harrower, and E. B. Dennis. 2022. rbms: Computing generalised abundance indices for butterfly monitoring count data. R package version 1.1.3.

Schmucki, R., G. Pe’er, D. B. Roy, C. Stefanescu, C. A. M. Van Swaay, T. H. Oliver, M. Kuussaari, A. J. Van Strien, L. Ries, J. Settele, M. Musche, J. Carnicer, O. Schweiger, T. M. Brereton, A. Harpke, J. Heliölä, E. Kühn, and R. Julliard. 2016. A regionally informed abundance index for supporting integrative analyses across butterfly monitoring schemes. Journal of Applied Ecology 53:501–510.

Settele, J., O. Kudrna, A. Harpke, I. Kühn, and C. Swaay, Van. 2008. Climatic risk atlas of European butterflies. Biodiversity and ecosystem risk assessment 1.

Shaw, M. R., C. Stefanescu, and S. Van Nouhuys. 2009. Parasitoids of European butterflies. Pages 130–156 in J. Settele, T. Shreeve, M. Konvicka, and H. Van Dyck, editors. Ecology of butterflies in Europe. Cambridge University Press, Cambridge.

Shorrocks, B., J. Rosewell, K. Edwards, and W. Atkinson. 1984. Interspecific competition is not a major organizing force in many insect communities. Nature 310:310–312.

Stefanescu, C., R. R. Askew, J. Corbera, and M. R. Shaw. 2012. Parasitism and migration in southern palaearctic populations of the painted lady butterfly, Vanessa cardui (Lepidoptera: Nymphalidae). European Journal of Entomology 109:85–94.

Stefanescu, C., P. Colom, J. M. Barea-Azcón, D. Horsfield, B. Komac, A. Miralles, M. R. Shaw, A. Ubach, and D. Gutiérrez. 2022. Larval parasitism in a specialist herbivore is explained by phenological synchrony and host plant availability. Journal of Animal Ecology 91:1010–1023.

Stiling, P. 1988. Density-Dependent Processes and Key Factors in Insect Populations. Journal of Animal Ecology 57:581–593.

Ubach, A., F. Páramo, M. Prohom, and C. Stefanescu. 2022. Weather and butterfly responses: a framework for understanding population dynamics in terms of species’ life-cycles and extreme climatic events. Oecologia 199:427–439.

Vigo, J., J. Carreras, A. Ferré, and J. J. Cambra, editors. 2005. Manual dels hàbitats de Catalunya: catàleg dels hàbitats naturals reconeguts en el territori català d’acord amb els criteris establerts pel “CORINE biotopes manual” de la Unió Europea. Departament de Medi Ambient i Habitatge.

Vila, R., C. Stefanescu, and J. M. Sesma. 2018. Guia de les papallones diürnes de Catalunya. Lynx Edici. Barcelona.

Whittaker, R. J., and J. M. Fernández-Palacios. 2007. Island biogeography: ecology, evolution, and conservation. Oxford University Press.

Yamamoto, N., J. Yokoyama, and M. Kawata. 2007. Relative resource abundance explains butterfly biodiversity in island communities. Proceedings of the National Academy of Sciences of the United States of America 104:10524–10529.

