## Supporting information for "Spatial variation in relative abundances of two butterfly species sharing both host plant and natural enemies"

**Tables**

**Table S1.** Results of the GLMMs testing the density dependent effects on the inter-annual population growth of the *Gonepteryx* species. Values of the coefficients for the variables included in the best-fit models based on AIC are shown. Values of the variables with a significant effect are represented in bold with asterisks (* *P* < 0.1; ** *P* < 0.05; *** *P* < 0.01).

| **Species** | **Cleopatra *(t-1)*** | **Brimstone**  ***(t-1)*** | ***Gonepteryx sp.***  ***(t-1)*** | **dAIC** |
| --- | --- | --- | --- | --- |
| Cleopatra (population growth) | **-0.5***** |  |  | 0 |
|  | **-0.5***** | -0.02 |  | 1.8 |
|  |  |  | **-0.45***** | 41 |
|  |  | **-0.07***** |  | 88.7 |
| Brimstone  (population growth) | **0.07*** | **-0.62***** |  | 0 |
|  |  | **-0.6***** |  | 1.4 |
|  |  |  | **-0.53***** | 118.9 |
|  | **-0.06***** |  |  | 132.8 |

**Table S2.** Comparison of density dependence effect by biogeographic/climatic region. Values of the coefficients are shown and significant differences between groups are represented in bold with asterisks (* *P* < 0.1; ** *P* < 0.05; *** *P* < 0.01).

| **Species** | **Climatic region**  **comparison** | **Brimstone**  ***(t-1)*** | **Cleopatra**  ***(t-1)*** |
| --- | --- | --- | --- |
| Brimstone (population growth) | Mesic vs xeric | **0.17 **** |  |
|  | Island vs Cat xeric | 0.09 |  |
| Cleopatra  (population growth) | Alpine vs mesic |  | **0.21**** |

**Figures**

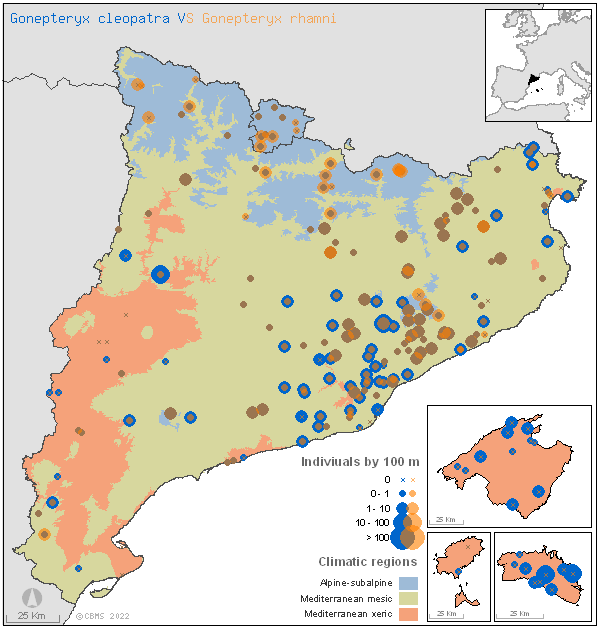

**Figure S1.** Annual relative abundance of Brimstone (in blue) and Cleopatra (in orange) in the study sites. Source: www.catalanbms.org

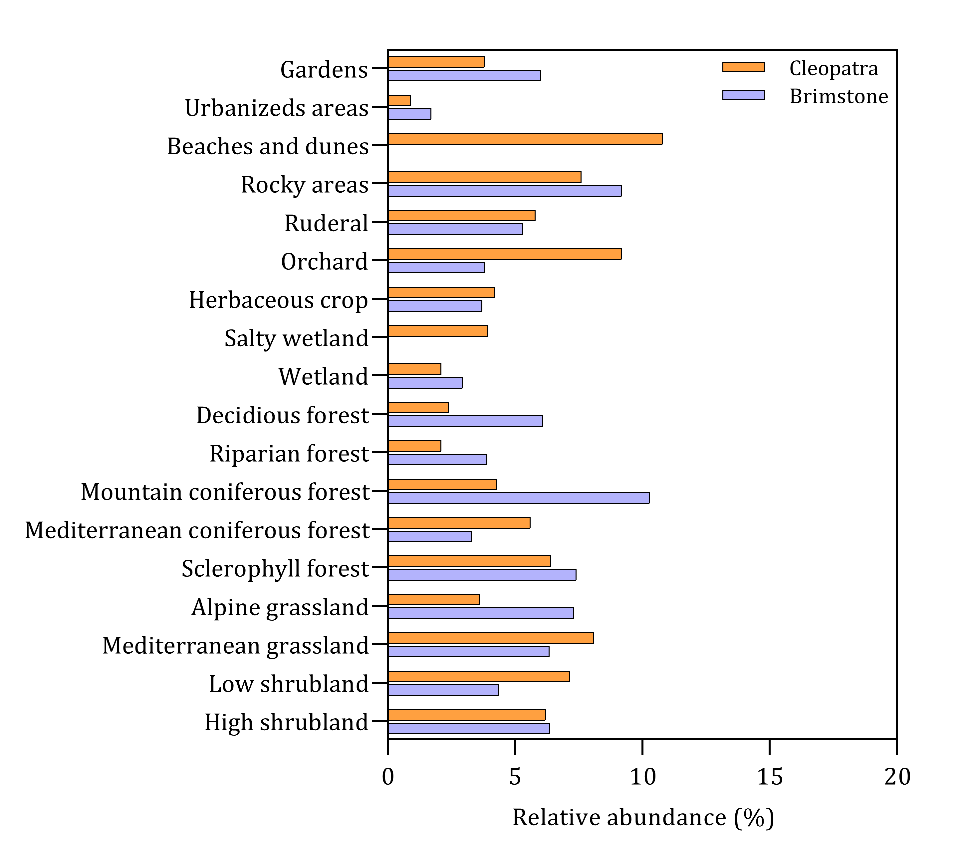

**Figure S2.** Relative abundances (%) of the adult butterflies for the habitats present in the study sites. Source: www.catalanbms.org

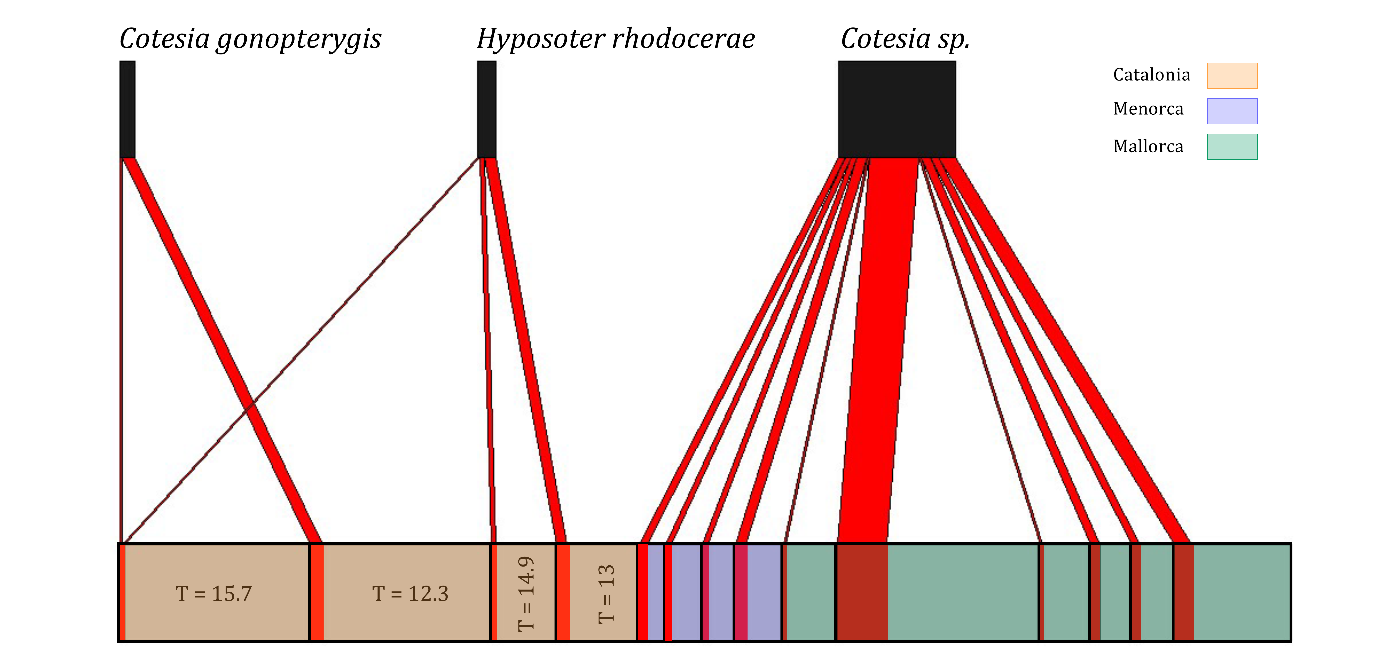

**Figure S3.** Bipartite plot (Dormann et al. 2009) showing quantitative associations between *Gonepteryx* larvae and its parasitoids in different study sites.The bottom boxes represent, for each study site, the number of larvae that were parasitized (red rectangle) and the number of larvae that developed into adults (coloured rectangle). For each species of parasitoid, the size of the box is proportional to its abundance in the samples (i.e. the number of larvae that were parasitized). Annual mean temperatures are indicated for the mainland sites in degree Celsius (T).

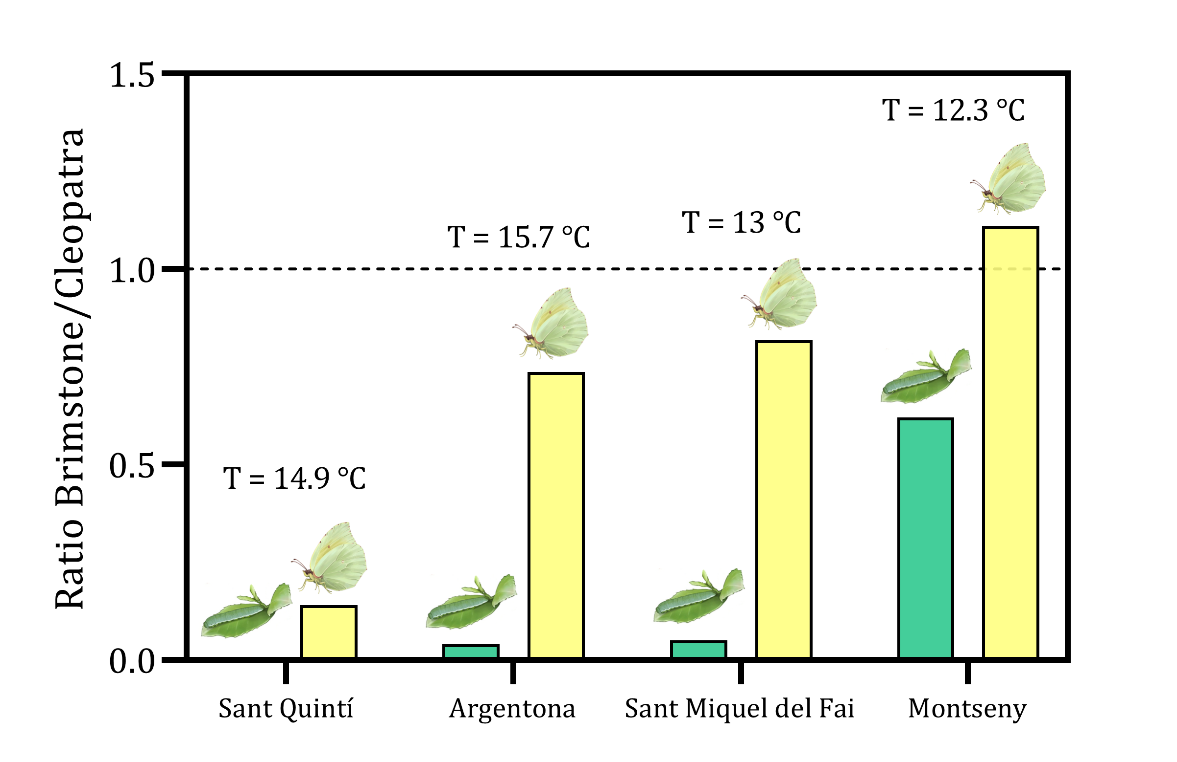

**Figure S4.** Ratio of number of individuals of Brimstone/Cleopatra in the mainland sites. Ratio for larvae individuals in green and ratio for adults in yellow. Annual mean temperatures for each site are indicated (T).
